# Rootstock effects on scion phenotypes in a ‘Chambourcin’ experimental vineyard

**DOI:** 10.1101/484212

**Authors:** Zoë Migicovsky, Zachary N. Harris, Laura L. Klein, Mao Li, Adam McDermaid, Daniel H. Chitwood, Anne Fennell, Laszlo G. Kovacs, Misha Kwasniewski, Jason P. Londo, Qin Ma, Allison J. Miller

## Abstract

Understanding how root systems modulate shoot system phenotypes is a fundamental question in plant biology and will be useful in developing resilient agricultural crops. Grafting is a common horticultural practice that joins the roots (rootstock) of one plant to the shoot (scion) of another, providing an excellent method for investigating how these two organ systems affect each other. In this study, we use the French-American hybrid grapevine ‘Chambourcin’ (*Vitis* L.) as a model to explore the rootstock-scion relationship. We examined leaf shape, ion concentrations, and gene expression in ‘Chambourcin’ grown own-rooted as well as grafted to three different rootstocks (‘SO4’, ‘1103P’ and ‘3309C’) across two years and three different irrigation treatments. Results described here demonstrate that 1) the largest source of variation in leaf shape stems from the interaction of rootstock by irrigation; 2) leaf position, but also rootstock and rootstock by irrigation interaction, are the primary sources of variation in leaf ion concentrations; and 3) gene expression in scion leaves exhibited significantly different patterns of gene expression from ungrafted vines, and these expression patterns were rootstock-specific. Our work provides an initial description of the subtle and complex effect of grafting on ‘Chambourcin’ leaf morphology, ionomics and gene expression in grapevine scions. Further work across multiple years, environments and additional phenotypes is required in order to determine how the relationship between the rootstock and the scion can best be leveraged for adapting grapevines to a changing climate.

## Introduction

Root and shoot systems operate in dramatically different environments and provide unique roles within a plant. These functionally distinct below- and above-ground parts are inextricably linked at the organismal level. Understanding the impact of roots on shoot system phenotypes, and conversely, how variation in the shoot influences the roots of a plant, are fundamental questions in plant biology. A further understanding of this interaction also has important agricultural implications, since selection for traits like root architecture and physiology can enhance stress tolerance and yield^1^.

In over 70 major crops, selection for root and shoot system traits have been decoupled through the process of grafting. Grafting is an ancient horticultural technique that creates a composite plant by surgically attaching the roots from one plant (the rootstock) to the shoot (the scion) of another, joining their vascular and cambial systems^2^. Grafting was originally implemented for easier clonal propagation, but today this method achieves a variety of agricultural goals, including drought tolerance, dwarfing, and disease resistance^1^. Beyond its practical implications, grafting offers an unique opportunity to independently manipulate parts of the plant to understand how roots impact shoots, and vice versa.

Grapevine (*Vitis* L. spp.) is an excellent model for examining rootstock-scion interactions due to the ease of cloning, available genomic resources, ability to grow across diverse environments, and high economic value. Widespread grafting of grapevine began in the late 19^th^ century after the European wine industry was devastated by the spread of phylloxera (*Daktulosphaira vitifoliae* Fitch), an aphid-like insect introduced from North America. While many North American *Vitis* species can withstand phylloxera infestations, roots of the European wine grape *Vitis vinifera* L. cannot tolerate phylloxera attacks, which lead to a rapid decline in vigour and often death^3^. However, *V. vinifera* vines with susceptible roots can be grafted to phylloxera-tolerant North American *Vitis* rootstocks, thus circumventing phylloxera sensitivity. Worldwide more than 80% of all vineyards grow vines grafted onto rootstocks composed of American *Vitis* species or hybrids^3^.

Although initial grapevine grafting was driven by the need for phylloxera tolerance, additional benefits exist. For example, certain *Vitis* rootstocks provide resistance to additional pests and pathogens such as nematodes^4^. Rootstocks can also be used to increase tolerance to abiotic stresses including drought^5,6^, salinity^7^, and calcareous soils^8^. Lastly, grafting can modify mineral nutrition^9^, scion vigor^10^, rate of ripening^11^, and fruit phenolic compounds^12^. Thus, grafting is a valuable tool for improving grapevine fruit quality and response to stress.

Grapevine producers rely on experimental trials to identify elite rootstocks that will best fit their specific growing conditions. Most commonly used grapevine rootstocks are hybrid derivatives of two or three phylloxera-tolerant native North American species, *Vitis riparia* and *Vitis rupestris*, which root easily from dormant cuttings, and *Vitis cinerea* var. *helleri* (*Vitis berlandieri*), which is adapted to chalky soils^13^. Interestingly, despite the global diversity of soils, climates and grape varieties, only a handful of rootstock cultivars derived from these three species are in widespread use^3^.

The result of over a century of grafting grapevines is a wealth of information characterizing graft-transmissible traits. In some cases, the biological mechanisms underlying beneficial effects are now understood. For example, salt (NaCl) tolerant rootstocks can exclude sodium (Na^+^) from the shoot, due to *VisHKT1;1*, a gene which can could serve as a valuable genetic marker for rootstock breeding^14^. However, for many other rootstock traits, the genetic underpinnings remain unknown. For example, the ability of a particular rootstock to protect the scion from iron deficiency was associated with an increase in root biomass along with a reduction of scion growth, but the molecular basis of this relationship is yet to be elucidated(Covarrubias et al 2016). Improved understanding of rootstock-scion interaction can enhance rootstock breeding for changing climates^15^ and evolving pest and pathogen pressures^13,16^.

While many facets of rootstock and scion interactions are still poorly understood, this study focuses on quantifying the effects of rootstocks on scion leaf shape, ion concentration, and gene expression. Traditionally, grapevine leaf morphology played a major role in the field of ampelography since it can be used to distinguish grapevine cultivars(Galet 1979). We examine the ability of quantitative measurements of leaf shape to discern subtle effects of rootstocks on scion development. We also examine the effect of rootstocks on leaf ionomic profiles, consisting of mineral nutrients and trace elements(Salt et al., 2008). Rootstocks, which limit or enhance the transport of particular elements, could facilitate grape-growing in regions with suboptimal soil conditions. Lastly, we examine patterns of gene expression between grafted and own-rooted vines. Recent work has described rootstock-induced differential gene expression in response to soil conditions such as nitrogen availability^17^. However, research so far has focused primarily on evaluating rootstocks with known contrasting effects under stressful conditions, and a broader understanding is still needed. Ultimately, understanding how a rootstock effects scion traits can further our understanding of root-shoot communication and provide insight when selecting parents or progeny in a rootstock breeding program.

To better understand the rootstock-scion relationship, our work examines ‘Chambourcin,’ a French-American hybrid grape of commercial importance ^18^. We examined ‘Chambourcin’ grown own-rooted as well as grafted onto three different rootstocks (‘SO4’, ‘1103P’ and ‘3309C’) across two years and three different irrigation treatments. Using comprehensive leaf shape analysis, ion concentration as determined by ICP-MS, and patterns of gene expression, we test the hypothesis that scion traits can be manipulated by different rootstock genotypes. We also examine the relative contribution of other experimental factors (e.g. irrigation treatment) as it relates to the potential for rootstock environment interactions to modulate scion phenotypes.

## Results

### Leaf shape

Using shape descriptors to examine variation in leaf morphology, we found that a significant amount of variance in aspect ratio (6.64%) and roundness (6.66%) measurements are explained by year of collection while variation in circularity is significantly explained by rootstock (5.00%) and irrigation factors (1.66%) (Figure 6A, Table S1). We visualized variation in the circularity based on irrigation (Figure 1B) and rootstock (Figure 1C), finding that leaves from vines which had full irrigation the year prior tended to have more subtle lobing and serration (i.e., higher circularity values). Circularity values were also higher for leaves of scions grafted to ‘1103P’ rootstocks compared to other rootstock treatments (Figure 1C). Lastly, a significant but minor amount of the variance in leaf solidity, which captures serrations or lobing, is explained by rootstock (2.35%) and rootstock by irrigation interaction (1.06%).

**Figure 1.**
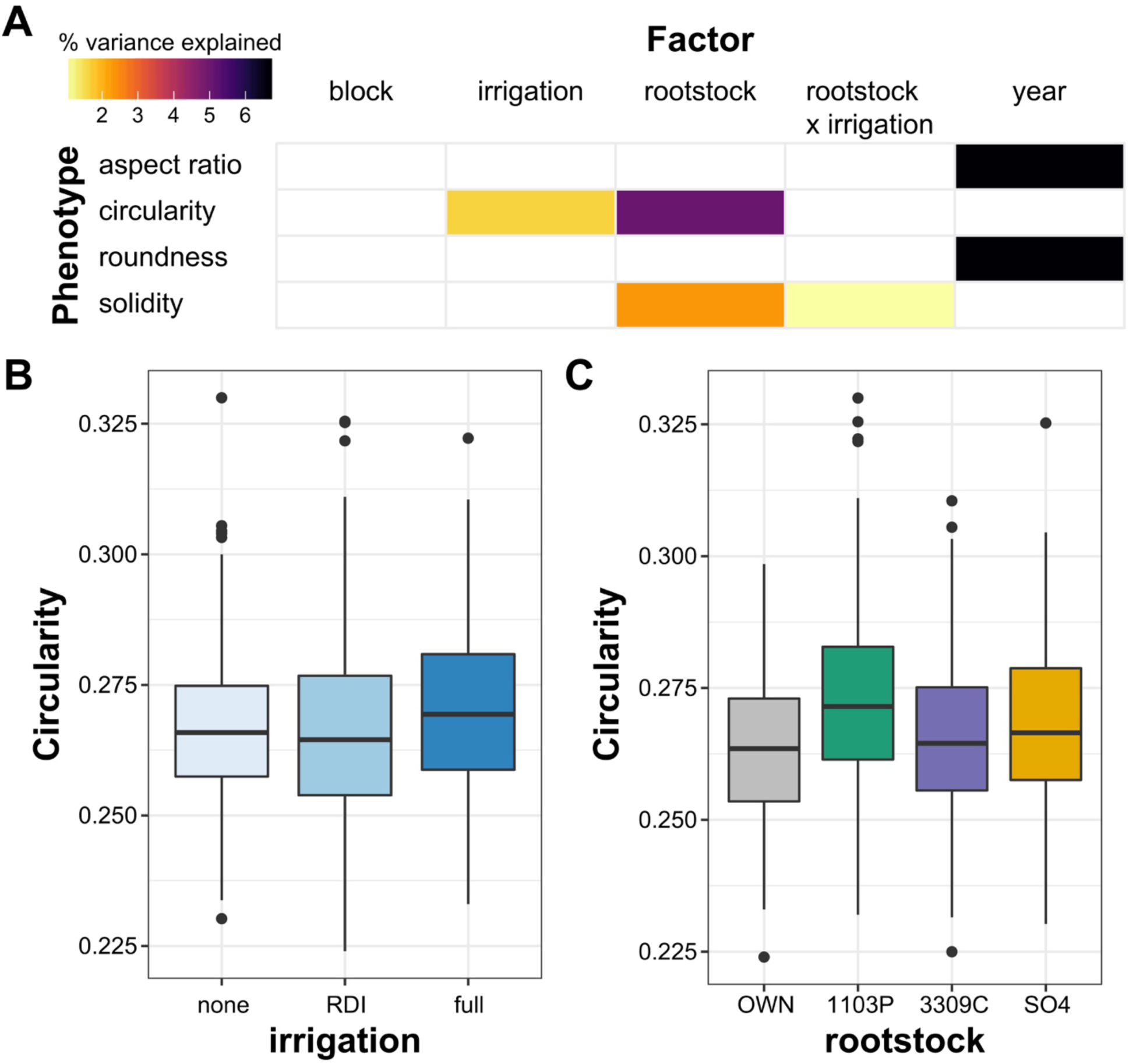
Variation in leaf morphology assessed using the shape descriptors aspect ratio, circularity, roundness and solidity. (A) A linear model was estimated for shape descriptors including the factors block, year, rootstock, irrigation and rootstock by irrigation. Only factors which explained a significant portion of the variance (p <0.05) are plotted. The percent variance explained by each factor is indicated using color. (B) Boxplots indicating circularity based on irrigation treatment. (C) Boxplots indicating circularity based on rootstock.

To examine the contours of grapevine leaf shape more comprehensively, we performed a persistent homology analysis, followed by PCA (Figure 2, Table S2). For PC1, which explains 17.56% of the variation in leaf shape, the primary source of variation described by our measurements is year (3.47%), followed by block (2.90%). However, across many morphometric PCs examined, the rootstock by irrigation interaction describes more variation than any other factor assessed. Of the 26 significant relationships (p<0.05) identified for PCs 1 to 20, 12 are for rootstock by irrigation interaction, followed by 5 for year. In contrast, rootstock explains a significant portion of the variation in leaf shape for 4 PCs, while irrigation is a significant factor for 2 PCs. Thus, changes in leaf shape measured using topological, persistent homology approach are most affected by the interaction of rootstock by irrigation, although year and block (which reflects position in the vineyard) are important as well.

**Figure 2.**
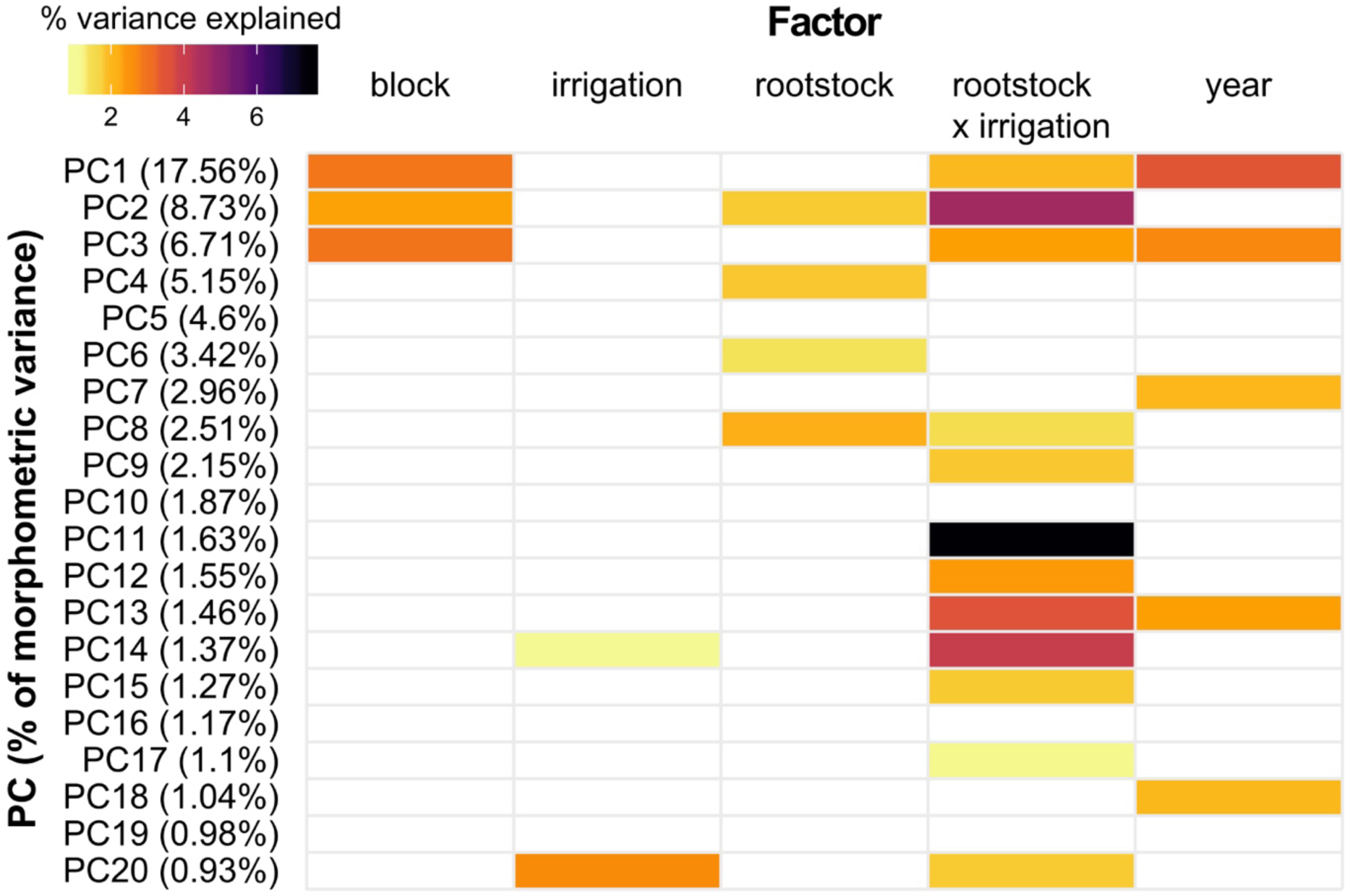
A linear model was estimated for morphometric PCs 1 to 20 including the factors block, year, rootstock, irrigation and rootstock by irrigation. The amount of variance explained by each PC is listed in parenthesis and the first 20 PCs capture a total of 68.13% of the variance in leaf shape. Only factors which explained a significant portion of the variance (p <0.05) are plotted. The percent variance explained by each factor is indicated using color.

### Ion concentrations

We used the same linear model approach to estimate which factors described the most variation in the 17 elements we examined for leaf ionomics (Figure 3, Table S3). In addition to the factors considered for leaf morphology, we assessed leaf position along the shoot (‘leaf’), a reflection of leaf developmental stage. As a result, our model identifies potential factors contributing to differences in ion concentrations including block, irrigation, irrigation by leaf interaction, leaf, rootstock, rootstock by irrigation interaction, rootstock by leaf interaction, and year as potential factors contributing to ionomic differences. The concentrations of ions in ‘Chambourcin’ leaves was most affected by leaf position, which explained a significant amount of the variance for 16 of the 17 elements we examined, ranging from 7.85% for nickel (Ni) to 60.89% for potassium (K). Over 50% of the variance in Calcium (Ca) can be explained using leaf position, and over 36% of the variance in manganese (Mn), aluminium (Al), and rubidium (Rb) can be explained. Rootstock also contributed to a substantial amount of variation in ion profile; it was a significant factor for 13 elements, most notably Ni, where it explained 24.94% of the variation. Lastly, the interaction between rootstock and irrigation was a significant factor for 17 elements, explaining over 30% of the variance for phosphorus (P), strontium (Sr), Rb, and molybdenum (Mo). In comparison, all other factors explained a maximum of 3.75% of the variation for any particular element.

**Figure 3.**
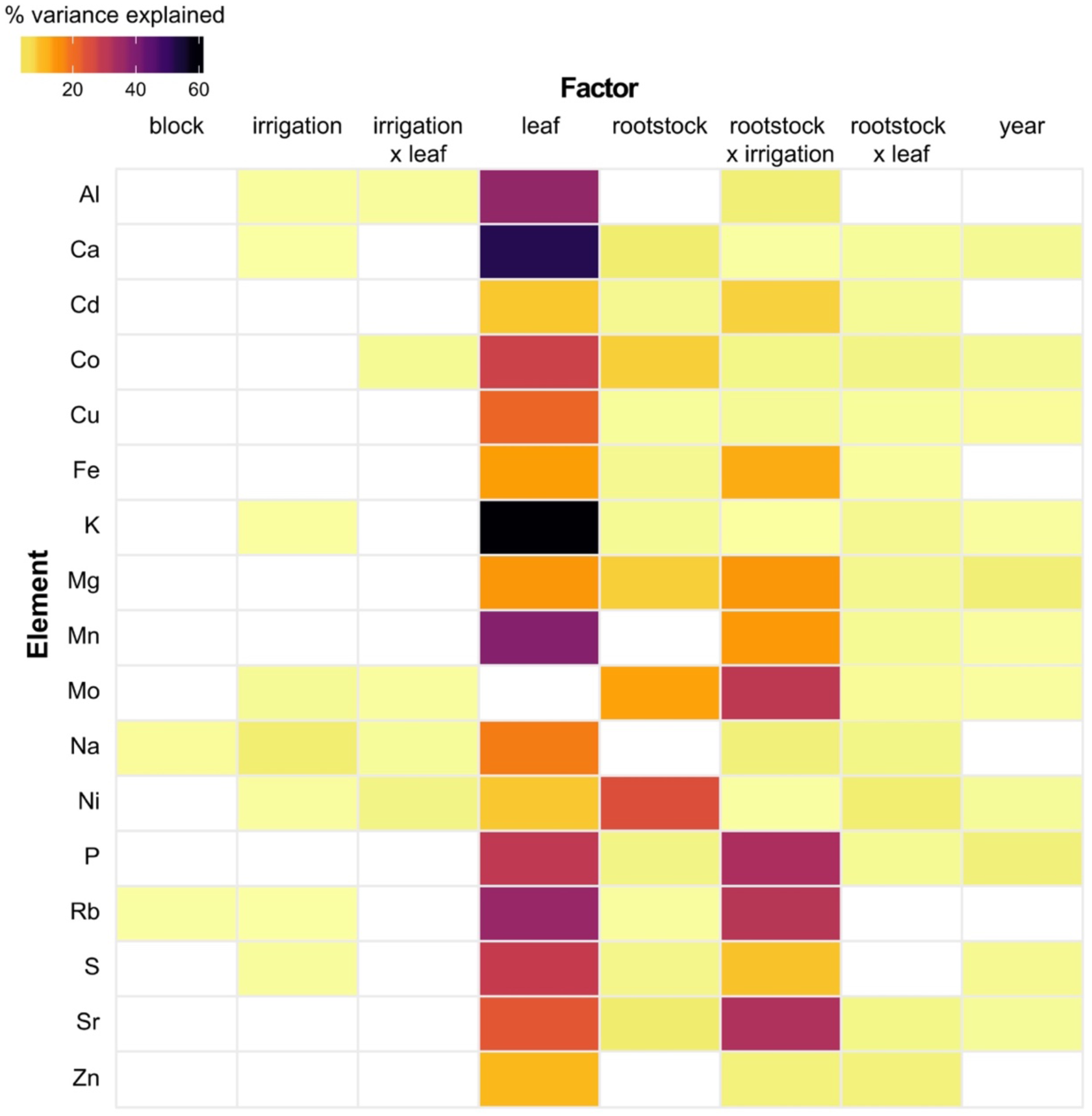
A linear model was estimated for each element including the factors block, year, rootstock, irrigation, rootstock × irrigation, leaf, rootstock × leaf, and irrigation by leaf. Only factors which explained a significant portion of the variance (p <0.05) are plotted. The percent variance explained by each factor is indicated using color.

By examining variation for each element across these factors of interest (Figure S2) we were able to observe several trends (Figure 4). For example, we found that Ca concentration increased in older ‘Chambourcin’ leaves (Figure 4A), while K concentration decreased in older leaves (Figure 4B). Across different rootstock treatments, the leaves of ‘Chambourcin’ grafted to ‘SO4’ generally had the highest concentration of Ni (Figure 4C). We also observed that Mo concentrations tended to increase from own-rooted, to ‘1103P’, to ‘3309C’, to ‘SO4’. Within a particular rootstock, vines which had been fully or partially irrigated the previous season tended to have ‘Chambourcin’ leaves with higher concentrations of Mo than those which had not been irrigated previously (Figure 4D).

**Figure 4.**
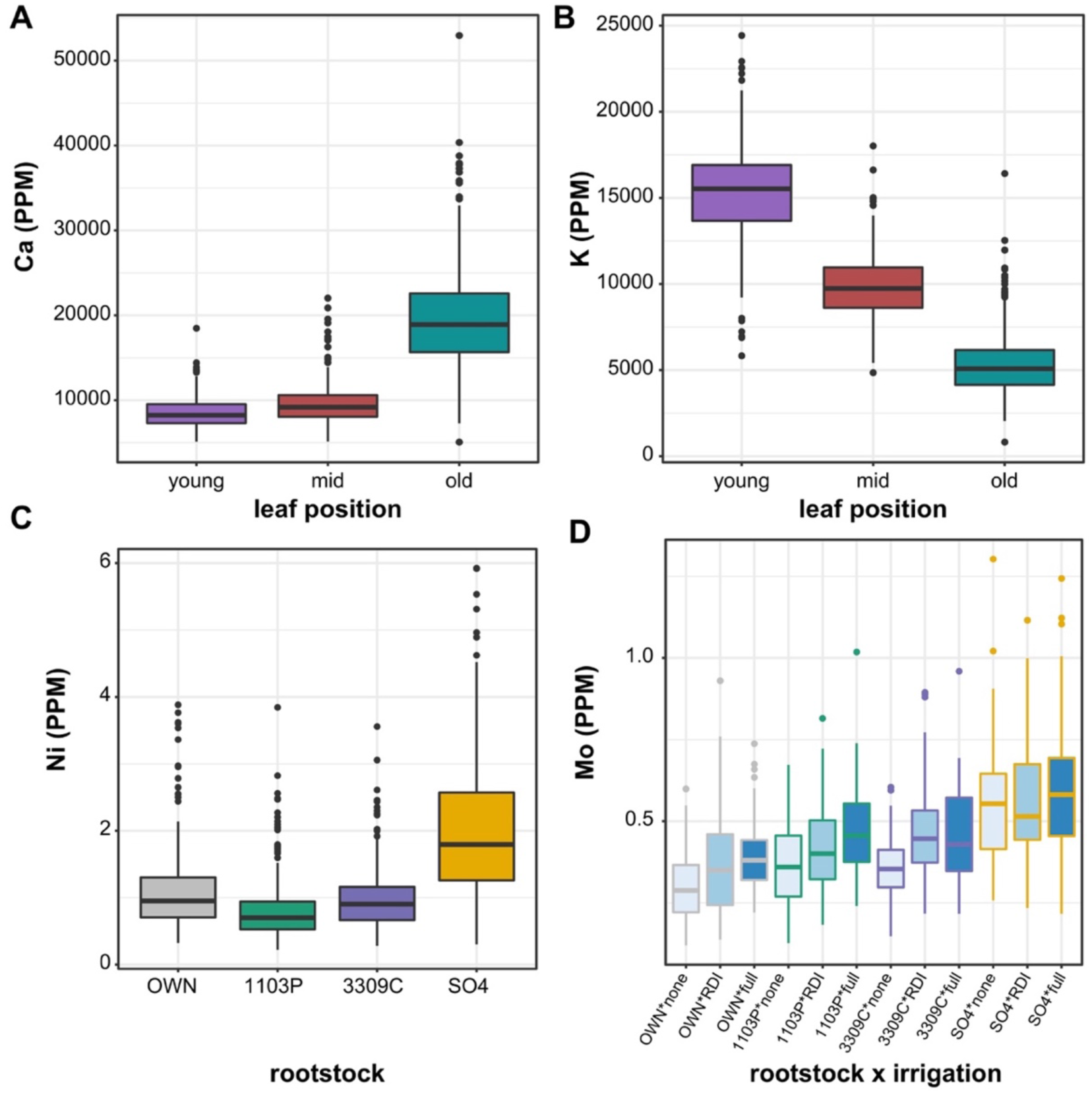
Boxplots showing the distribution of elements by on the factor that explained the largest amount of variance. Distributions shown are: (A) Ca due to leaf position (B) K due to leaf position (C) Ni due to rootstock (D) Mo due to rootstock by irrigation interaction.

### Gene expression

We used all gene expression RPKM values to test for positively enriched VitisNet Pathways by comparing ‘Chambourcin’ grafted to each individual rootstock with own-rooted ‘Chambourcin’ vines. Each rootstock has 8 unique enriched pathways. The pathways enriched in ‘1103P’ include circadian rhythm and phenylalanine metabolism. We combined all grafted ‘Chambourcin’ and compared them to own-rooted vines to determine the impact of grafting, identifying 17 enriched pathways in grafted vines. All pathways are listed in Table S4.

Next, we used a regression fit that accounted for replicate, block, and rootstock for each gene significantly expressed in own-rooted ‘Chambourcin’ vines to determine which genes had differing patterns of expression in ‘Chambourcin’ scions when grafted (Figure 5). In total, there were 513 genes in own-rooted ‘Chambourcin’ vines with significant expression. Of these genes, 121 were not significantly differentially expressed in any of the rootstock treatments, after accounting for block, which represented both spatial and temporal variation. Comparing grafted vines to own-rooted vines, 5 genes exhibited significantly different expression profiles in all three grafted vines compared to own-rooted vines. The only annotated gene among these five is an isoamylase protein. Relative to own-rooted vines, there were 105 genes which had significantly different expression patterns only in ‘Chambourcin grafted to ‘3309C’, 96 which differed only in ‘Chambourcin’ grafted to ‘1103P’, and 89 which differed only in ‘Chambourcin’ grafted to ‘SO4’ (Table S5; Table S6). Pathway enrichment analysis was used to examine these rootstock specific genes. While no major enrichment was observed for the ‘3309C’ and ‘SO4’ genes, ‘1103P’ vines had a significant number of genes involved in phenylalanine metabolism (4 DEGs, p = 4.84 × 10^−6^) and auxin biosynthesis (3 DEGs, p = 1.74 x10^−5^) pathways (Table S6).

**Figure 5.**
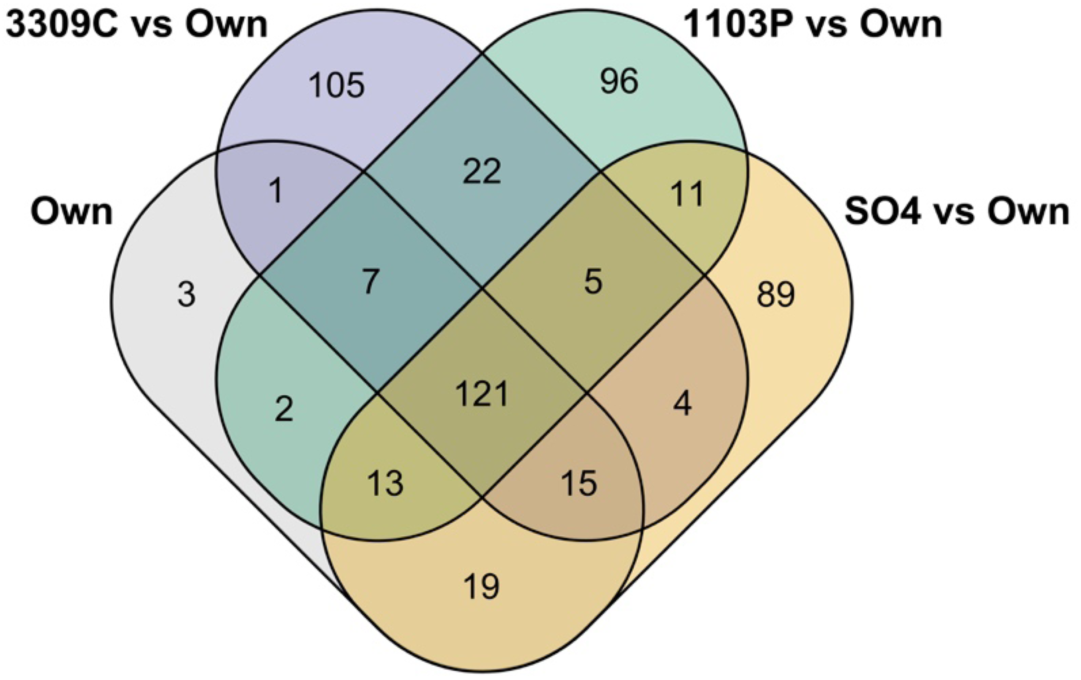
All genes significantly expressed in own-rooted vines as determined using a regression fit which considered block, replicate and rootstock, were compared to each rootstock. The Venn diagram indicates the number of genes which were significantly differentially expressed when a particular rootstock was compared to own-rooted vines.

## Discussion

Grafting offers an excellent opportunity to understand how roots modulate scion phenotypes through the experimental manipulation of root systems and grafted scions. Our study uses grapevine as a model to quantify the effect of rootstock on leaf shape, ion concentrations, and gene expression in the scion. Results described here demonstrate that genetically distinct root systems interact in unique ways with seasonal water availability to influence shoot system phenotypes in grafted plants.

### Leaf shape is modulated by the interaction of rootstock and irrigation

The grapevine genus is well-known for extensive within and among-species variation in leaf shape^19,20^. Previous work demonstrated that the genetic underpinnings of leaf shape are evolutionarily conserved within species, while developmental constraints and environmental influences such as light, temperature, and water availability affect leaf shape variation among genotypes and within individuals^21–23^. We collected leaves from approximately the same developmental stage (i.e. position on the shoot) from vines of ‘Chambourcin’ to minimize leaf shape differences due to position along the vine (i.e., heteroblasty^24^).

We measured leaf shape using two approaches: shape descriptors, a common digital morphometric technique that captures simple shape differences, and persistent homology, a comprehensive morphometric technique, which allowed us to detect complex and subtle variation in shape. We observe interannual variation in leaf shape using both shape descriptors and persistent homology. Of the two methods, shape descriptors capture variation across years, but generally do not vary due to rootstock by irrigation effects. For example, approximately 1% of variation in the solidity measurement was significantly explained by rootstock by irrigation, while the same interaction effect was a significant factor for 12 of the 20 morphometric PCs examined, explaining up to 7.53% of the variation for a particular PC. This reflects the ability of digital morphometric techniques to detect subtle, significant statistical effects on leaf shape in a targeted way: unlike the persistent homology approach, statistical differences in solidity correspond to serration and lobing, suggesting these features vary across years.

In contrast, persistent homology was able to detect a significant portion of morphological variation in leaf shape due to position in the vineyard or block. Persistent homology uses a comprehensive method for quantifying shape, and likely picks up on intricate leaf shape differences that traditional methods miss. With this method, we were able to demonstrate that the rootstock interacting with irrigation effect shifts the shape of ‘Chambourcin’ leaves in comprehensive, detectable ways. Recent work in apple described a heritable basis for leaf shape, as described using persistent homology^25^. Our work suggests that rootstocks could be used to modulate variation in leaf shape in the scion, especially under varying environmental conditions such as access to water/irrigation treatments. More importantly, our results suggest that rootstocks can modulate scion development and patterning, that signals from the root (whether molecular or physiological in nature) affect patterning within scion meristems. Although some molecular evidence supports such long-distance coordination of developmental patterning^26^, its prevalence and manifestation across plants remains understudied, even though it is critical to understand as rootstocks are used more widely to modulate scion phenotypes.

In addition to our work, other studies in grapevine have identified scion leaf shape modulation under different rootstock and irrigation treatments. Tsialtas et al. (2008) examined ‘Cabernet-Sauvignon’ grafted on ‘1103P’ and ‘SO4’ rootstocks under 3 different irrigation treatments at 3 time points (bunch closure, veraison and ripeness). The work found that while rootstock, irrigation and rootstock by irrigation did not have a significant effect on leaf morphology, the rootstock by irrigation by time interaction was significant for all leaf shape measurements assessed^27^. In addition, recent work evaluating the leaves of ‘Italia’ grapes grown own-rooted and grafted to 2 rootstocks under 2 irrigation conditions, found that leaf area was significantly affected by rootstock by irrigation interaction^28^. Thus, it is clear that the influence of rootstock on leaf shape is a complicated relationship that is at least partially influenced by other factors including irrigation. Pairing these data with physiology and ionomics may help identify more precisely the effect of rootstock by irrigation on leaf shape in future studies.

### Scion elemental composition is mostly affected by leaf position, but also rootstock and rootstock by irrigation interaction effects

The interaction between root system and elemental composition in grapevine shoot systems has been an area of great research interest in viticulture^9,29^. The grapevine industry places enormous importance on *terroir*, the physical environment in which a grapevine is grown, to determine the sensory experience and economic value of wine^30^. Indeed, research shows that available soil nutrients can be transported and stored in different plant tissues^31^ and that rootstock can affect different ion uptake^32^. The ability of the rootstock to impact ion uptake in grapevine is of particular note because such differences can have a pronounced effect on wine quality. Soil elements such as Mg, Mn, and Mo are present in berries throughout wine production (i.e., harvest to bottling), depending on the concentration of these elements in a given geographic region^30^. Our study builds upon a body of literature that demonstrates rootstock selection modulates the movement and concentration of elements in scion tissues^9,33^.

In our work, the position of the leaf on the shoot (the developmental stage of the leaf) explains the largest amount of variation observed in most ions. Previous work by Huber et al.^34^ found that position along the main stem had a profound effect on seed composition in soybean. We examined 17 elements and found that for 13 the primary source of variation explaining ion concentration was leaf position. New leaves must rely transpiration to transport Ca from the xylem, and since transpiration is low in young leaves, we observe that younger leaves had lower concentrations of Ca than older leaves^35^. Al and Mn also decreased in younger leaves, while K and Rb increased. These elements provide examples of the changes that occur in elemental composition as leaves develop and age, regardless of rootstock.

While the primary source of variation in ion concentrations was leaf position, a significant amount of variation was explained by the interaction between rootstock and irrigation for all 17 elements, while rootstock explained a significant amount of variation for 13 elements. Either rootstock or rootstock by irrigation also explained >10% of the variation for Fe, Mg, Mn, Mo, Ni, P, Rb and Sr. Previous work identified that grafting ‘Négette’ vines onto ‘SO4’ resulted in higher K and lower Ca and Mg concentrations compared to ‘3309C’ and ‘101-14 Mgt’^36^. While we did not detect a similar pattern in the leaves of ‘Chambourcin’ scions, we found that vines grafted to ‘SO4’ had higher concentrations of Ni than vines grown own-rooted or grafted to ‘3309C’ or ‘1103P’. Across the United States, Ni is highest in serpentine soil areas of California^37^. Serpentine soil increases Ni accumulation in grapevine roots, with previous work also finding a significant positive correlation between Ni in the soil and leaves. However, the transfer of Ni from grapevine roots to grapes was low^38^. While further testing in serpentine soil is still required, our work provides evidence that ‘SO4’ may not be an optimal rootstock choice for high Ni soils, since excess Ni may cause toxicity limiting crop production^39^.

In contrast to leaf position, rootstock, and the interaction of rootstock and irrigation, we generally do not see a significant effect of irrigation on ion concentrations. However, our samples were collected prior to the start of irrigation treatments in 2014 and 2016, and thus, any response to irrigation would be due to historical conditions and chronic stress, rather than current, acute stress. Future work sampling throughout the growing season, both before and after the initiation of irrigation treatments, will be required to assess how historical and current water conditions influence ion concentrations.

Beyond assessing variation in each element independently, previous work has demonstrated that elements interact with each other^40^. Consequently, it is not surprising that we find so many elements influenced by the same factor. In fact, leaf position, rootstock, and rootstock by irrigation interaction each explain a significant amount of variation in at least 13 of the 17 elements, and this broad effect may indicate interaction between elements. It is clear that the root system, the environment – including irrigation and position within the shoot – and the interaction between the two are critical in determining ion concentrations, and a further understanding of these complex relationships is still necessary.

### Rootstocks alter scion gene expression

Grafting alters scion phenotypes by affecting the availability of water and nutrients, changes which may contribute to modified patterns of gene expression in the scion. Rootstock modulation of scion phenotypes is evident in stressful conditions, as has been demonstrated in many major crops including tomato, apple, citrus, and grapevine, among others^41–44^. However, basic differences in gene expression in grafted plants relative to ungrafted individuals remain underexplored^45^. Thus, grafting to a common scion provides an excellent opportunity to better understand how environment impacts shoot system phenotypes in plants under normal growing conditions.

In our study, we assessed the influence of root systems on gene expression in shoot systems by contrasting gene expression in ‘Chambourcin’ grafted to three different rootstocks relative to own-rooted vines. When comparing DEGs expressed in grafted vines to own-rooted vines, we found a similar number of genes (89-105) which were only differentially expressed in one rootstock treatment. This relatively low number of genes may indicate that variation in the scion transcriptome is predominantly under local genotype (scion) control and not dependant on signalling from the rootstock. Given the life history of grapevine, a liana with typically long distances between roots and shoots, it is perhaps not a surprising result. Only five genes were consistent in their patterns of differential expression across all rootstocks when compared to own-rooted vines, indicating that there are rootstock-specific effects on scion gene expression.

We also examined the influence of grafting to different rootstocks on specific pathways using two methods: first, by using all expressed genes to assess pathway enrichment, and second, by only including genes determined to be significantly differentially expressed. Prior to the inclusion of block in our analysis, the pathway analysis detected enrichment of the circadian rhythm pathway in ‘1103P’ relative to own-rooted vines. Thus, even within a timespan of sampling (approximately 8 hours) it is necessary to consider the impact of time on changes in gene expression, and future work is needed to describe whether the impact of sampling time is rootstock-specific. Both techniques found unique pathways enriched in each rootstock, relative to own-rooted vines, providing further evidence that the effect of grafting on gene-expression is rootstock-specific.

Among the 96 genes with expression patterns that differed only between ‘1103P’ and own-rooted vines, both pathway analyses revealed an enrichment of those involved in phenylalanine metabolism, while only the analysis of DEGs showed enrichment for auxin biosynthesis. Although our work examined leaf tissue, these results are supported by previous work comparing ‘Cabernet Sauvignon’ grafted to ‘1103P’ and ‘M4’ rootstocks which found that genes involved in auxin action were one of the main categories with a rootstock effect in the berry, especially for skin tissue^46^. Most work examining rootstock effects on scion gene expression focuses on variation under conditions of stress such as iron chlorosis^47,48^. In comparison, our work examined the effect of multiple rootstocks under neutral environmental conditions, and this difference likely explains the subtle but quantifiable effect of rootstock on scion gene expression described here. Ultimately, we find that the graft-transmissible effects on a common scion are rootstock-specific. Further, our work also indicates that time of sampling may play a significant role in rootstock effects, and further work is needed to explore this complex interaction.

## Conclusions

Our work provides an initial description of the subtle and complex effect of grafting on leaf morphology, ionomics and gene expression in grapevine scions. We find that specific rootstocks have a distinct effect on many of the phenotypes, often interacting with the environment due to previous water availability. Leaf position in the shoot and block position in the vineyard, also strongly influenced phenotypic variation. Further work across multiple years and environments is required in order to determine how the relationship between the rootstock and the scion can best be leveraged for adapting grapevines to a changing climate.

## Materials and methods

### Study design and sampling

A ‘Chambourcin’ experimental vineyard was established in 2009 at The University of Missouri Southwest Center Agricultural Experiment Station in Mount Vernon, Missouri, USA. The vineyard includes own-rooted, ungrafted ‘Chambourcin’ vines as well as ‘Chambourcin’ vines grafted to three different rootstocks (‘3309C’ - *V. riparia* × *V. rupestris*; ‘1103P’ - *V. berlandieri* × *V. rupestris*; ‘SO4’ - *V. berlandieri* × *V. riparia*). The full factorial experiment with varied rootstock and irrigation regimes contains 288 vines: eight replicates of four root-scion combinations × nine vineyard rows with one of three irrigation treatments. The three irrigation regimes are: full replacement of evapotranspiration losses (ET), 50% replacement of ET, and non-irrigated, each replicated three times (Figure S1). Further description of the vineyard design is available in Maimaitiyiming et al., 2017^49^. Irrigation treatments began in 2013, with all vines receiving full irrigation during establishment. Irrigation treatments were initiated several weeks before veraison. Sampling of leaf tissue for morphometric and ionomic analyses occurred on June 18, 2014 and June 14, 2016, while tissue for gene expression analyses was sampled only on June 14, 2016. In both years, sampling occurred prior to the start of irrigation treatments, and thus, any effect of irrigation we observe is due to treatment from the previous year(s), when the buds/leaves/flower of the study years are formed. Data and code for this manuscript are available in a GitHub repository^50^.

### Leaf shape analyses

For leaf shape analyses, the middle four leaves from a single shoot were collected from each vine. Leaves were flattened, stored in plastic bags in coolers in the field, and transferred to a cold room in the lab. Within a few days of collection leaves were imaged using a Canon DS50000 document scanner. Leaves with margin damage were removed from analysis. The resulting dataset included 277 vines with 4 leaves and 6 vines with 2 leaves in 2014, and 284 vines with 4 leaves, and 2 vines with 2 leaves, in 2016.

Leaf scans were converted to binary (black and white) images in Matlab and then analyzed in ImageJ(Abrámoff et al., 2004) using shape descriptors including aspect ratio, circularity, roundness, and solidity, each of which captures a ratio describing variation in lobing and shape^51^. Shape descriptors were averaged across leaves from each plant. We performed linear modeling using the lm() function in R, accounting for variation in block (which reflects vineyard position), irrigation, rootstock, rootstock by irrigation interaction, and year. The percent variance explained by each factor was calculated using the anova() function, and only those with a significant p-value (<0.05) were visualized using the ggplot2 package in R (Wickham 2009).

To comprehensively measure leaf shape, we used a persistent homology approach, a type of Topological Data Analysis (TDA), to capture the outline of the leaf^52^. Each leaf was considered as a point cloud in which each pixel is a point (Figure 6A). A Gaussian density estimator was applied to each pixel reflecting the density of neighboring pixels (Figure 6B). In the context of leaf shape, pixels in serrations or lobes tend to have more neighbors than pixels that lie on relatively straight edges. 16 concentric annuli emanating from the centroid were multiplied by the Gaussian density estimator isolating subsets of the data (Figure 6C-F); the subsetted data (the rings) are arbitrary and are intended to provide an increased number of comparable topological features between leaves. Each ring is effectively a different set of topological features analyzed (that is, a set of 16 shapes for each grapevine leaf). In Figure 6G, high (red) and low (blue) values of the Gaussian density filtration function are visualized directly on a grapevine leaf shape. The number of connected components are monitored. As the filtration function is passed through (red-to-blue in Figure 6G), connected components will arise or merge with each other. Changes in number of connected components are the result of the position in the filtration function, which is the x-axis of the Euler characteristic curve (Figure 6G) and monitors the number of connected components (y-axis) as a function of the filtration. The Euler characteristic curves (one for each of 16 rings) were discretized. Further details of the analysis were previously published^53,54^. Binary images and persistent homology values are available for download^55^.

Persistent homology values were averaged across leaves for each plant and principal component analysis (PCA) was performed. The first 20 principal components (PCs) explained 68.13% of the total variance, and thus only these were included in downstream analyses. The morphometric PCs were included in a linear model which accounted for variation in rootstock, irrigation (which reflects historical treatment conditions) rootstock by irrigation interaction, year, and block. Lastly, we calculated how much of the total variance was explained by each trait, and factors explaining a significant portion of the variance (p<0.05) were visualized using the ggplot2 package in R(Wickham 2009).

**Figure 6.**
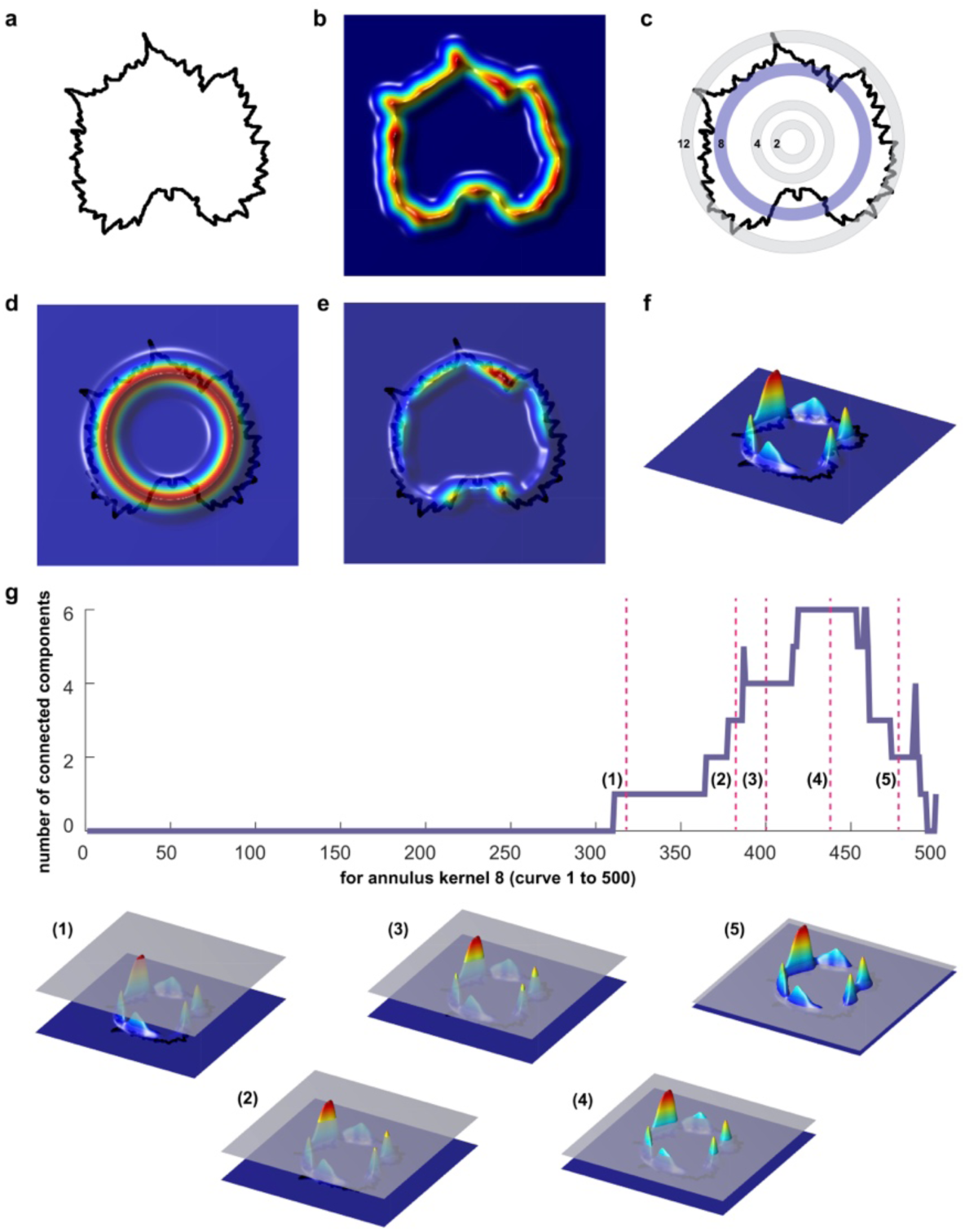
Quantifying leaf shape using persistent homology, a Topological Data Analysis (TDA) method. (A) A 2D point cloud represents each leaf contour. (B) A Gaussian density estimator estimates the density of neighboring pixels around each pixel. Pixels near serrations and lobes tend to have higher density values. (C) 16 concentric rings are used to partition the data as an (D) annulus kernel. (E) Multiplication of the annulus kernel by the Gaussian density estimator isolates sub-features of the leaf. (F) A side projection shows clearly the isolated density features of the leaf. (G) Proceeding from high density values to low (1-5) the number of connected components (a topological feature) is recorded as a function of density. The resulting curves from each ring are discretized and quantify leaf shape.

### Leaf ion concentration analyses

To investigate ion concentrations in the leaves, three leaves from different developmental stages were collected from a single shoot from each vine. The first leaf sampled came from the first node at the base of the shoot and was the oldest leaf on the shoot. The second leaf sampled (also used for morphometric analyses) came from the middle of the shoot. The third leaf was sampled at the tip of the shoot.

Each sample was processed for ionomic analysis at the Donald Danforth Plant Science Center (St. Louis, MO), as described in Ziegler et al.^56^, including correction for internal standards and standardization based on sample weight using custom scripts, with one minor modification in the dilution method. Samples were digested in 2.5mL nitric acid and then diluted to 10mL with ultrapure water. Instead of a second manual dilution, an ESI prepFAST autodiluter diluted samples an additional 5x inline with ultrapure water. The 2014 samples were analyzed using a Perkin Elmer Elan 6000 DRC-e ICP-MS run in standard mode. The 2016 samples were run with a Perkin Elmer NexION 350D ICP-MS with helium mode enabled. The standard used for normalizing samples in 2014 was rerun in December 2017 and all values from 2016 were adjusted to account for variation between instruments. The elements boron (B), selenium (Se) and arsenic (As) did not measure well in at least one year and were subsequently removed from the analysis for both years, resulting in 17 elements for subsequent analysis.

For both 2014 and 2016 ionomics data, we removed extreme outliers for each element with values less than 0.25 quantile − interquartile range*5, or greater than 0.75 quantile + interquartile range*5. After outlier removal, 703-794 samples per element remained for 2014 and 846 samples for 2016 remained. All samples were included in a linear model accounting for leaf, rootstock, irrigation, block, year, rootstock by irrigation interaction, rootstock by leaf interaction, and irrigation by leaf interaction, using the lm() function in R. Since tissue sampling occurred in June prior to the initiation of irrigation treatments, the effect of irrigation describes historical water conditions. The percent variance explained by each factor was calculated, and only those with a significant p-value (<0.05) were visualized.

### Gene expression analyses

We used RNA-seq to assess changes in gene expression in leaves of grafted and ungrafted ‘Chambourcin’ vines. Samples were collected from two rows with no irrigation treatment (rows 13 and 15, Figure S1) on June 14, 2016. Each row was composed of two blocks of vines, and within each block, we sampled two clonal replicates from each rootstock-scion combination, for a total of 32 samples. Samples were collected from row 15 column A to column H, and then from row 13 column A to column H. For each vine, we collected the first leaf at the tip of the shoot that was fully open (~16 mm in diameter). Leaf tissue was immediately flash frozen in liquid nitrogen and transported on dry ice before transferring to a -80°C freezer for storage.

Total RNA was extracted at the US Department of Agriculture Grape Genetics Research Unit (Geneva, NY) using standard extraction protocols for the Sigma Spectrum Plant RNA kit (Sigma Aldrich, Inc. St. Louis MO) with the following modification; addition of 3% w/v PVP40 added to the lysis buffer. Library construction was performed by Cofactor Genomics (http://cofactorgenomics.com, St. Louis, MO). Total RNA was incubated with mRNA capture beads in order to remove contaminating ribosomal RNA from the sample. The resulting poly(A)-captured mRNA was fragmented. First-strand cDNA synthesis was performed using reverse transcriptase and random primers in the presence of Actinomycin D, followed by second-strand cDNA synthesis with DNA polymerase I and RNase H. Double-stranded cDNA was end-repaired and A-tailed for subsequent adaptor ligation. Indexed adaptors were ligated to the A-tailed cDNA. Enrichment by PCR was performed to generate the final cDNA sequencing library. Libraries were sequenced as single-end 75 base pair reads on an Illumina NextSeq500 following the manufacturer’s protocols. The RNA-sequencing data have been uploaded to the NCBI Sequence Read Archive under BioProject PRJNA507625: SRA Accessions SRR8263050 - SRR8263077.

All samples were quality checked using FastQC v0.11.3(Andrews 2015). Reads were aligned to the 12Xv2 reference genome and the VCost.v3(Canaguier et al. 2017) reference annotation using HISAT2 v2.1.0(Kim et al. 2015). Counts were derived from the alignment with HTSeq^57^. Differential gene expression analysis was performed using the R package DESeq2^58^. After determining differential expression, the raw read counts were normalized using the DESeq2 normalization method of dividing each count by the size factors.

As an initial survey of the potential impact of rootstocks on gene expression, we conducted a Gene Set Enrichment Analysis (GSEA) using GSEA-P 2.0 (http://www.broad.mit.edu/GSEA) and 203 VitisNet pathways including at least 10 genes^59–63^. Enrichment was tested using normalized expression data (RPKM) for all genes, for each rootstock. The gene expression from leaf tissue (Chambourcin scion) for each root stock was compared separately to own-rooted ‘Chambourcin’ leaf gene expression, as well as, comparing all scion/rootstock combination gene expression to own-rooted leaves. For each comparison, we determined which pathways were up-regulated in grafted vines using GSEA. The GSEA-P 2.0 default parameters of 1000 permutations, nominal p-value (p < 0.05) and false discovery rate (FDR) q-value (q<0.25) were used to identify positive significantly enriched molecular pathways^60^.

Next, we determined significantly differentially expressed genes (DEGs) by comparing grafted vines to own-rooted vines. Samples from each block were collected chronologically, and thus, each block represented spatial variation as well as a particular time point. We performed a regression fit for each gene accounting for considering all variables (block, replicate, and rootstock) using the MaSigPro R package^64^. Using the p.vector() function, we returned a list of FDR-corrected significant genes, which were input into the T.fit() function, to perform stepwise regression, selecting the best regression model for each gene. The get.siggenes() function with the ‘vars=“groups”’ option was used to generate a list of genes with significant expression in own-rooted vines. Expression patterns for each rootstock were then compared to patterns in own-rooted vines, in order to determine which genes had significantly different expression profiles in a particular rootstock. Next, we used the suma2Venn() function to visualize overlap across rootstocks and own-rooted vines.

Lastly, we queried DEGs identified in each grafted ‘Chambourcin’ relative to own-rooted vines for statistical enrichment of metabolic and regulatory pathways, to determine if rootstock impacted specific aspects of vine biology. Unlike the initial GSEA assessment which included all genes, this analysis only included DEGs. We tested DEGs for pathway enrichment using the Vitisnet database^62^ and the VitisPathways tool^65^ using 100 permutations, a Fisher’s exact test of p< 0.05 and a permuted p value of p<0.05.

### Availability of data

Binary images and persistent homology values are available for download^55^. The RNA-sequencing data have been uploaded to the NCBI Sequence Read Archive under BioProject PRJNA507625: SRA Accessions SRR8263050 - SRR8263077. Data and code for this manuscript are available in a GitHub repository^50^.

## Supporting information

## Acknowledgements

R. Keith Striegler designed and established the ‘Chambourcin’ experimental vineyard at the University of Missouri Southwest Research Farm. We thank Greg Ziegler and the Baxter Laboratory (USDA-ARS/Danforth Center Ionomics Facility) for performing for ionomics work described in this study. We thank Margaret Frank (Cornell University), Viktoriya Coneva (Kenyon College), Rebekah Mohn (Donald Danforth Plant Science Center), Halley Fowler (Donald Danforth Plant Science Center), Stephanie Theiss (Donald Danforth Plant Science Center), and Alex Linan (Saint Louis University) for sampling leaves used for analysis.

This work was supported by Missouri Grape and Wine Institute, National Science Foundation Plant Genome Research Program 1546869, and Saint Louis University. This work was partially supported by appropriated funds to USDA-ARS-GGRU for project 8060-21220-006-00D. This project was also supported by the USDA National Institute of Food and Agriculture, and by Michigan State University AgBioResearch. We acknowledge support from National Science Foundation (NSF) Plant Genome Research Program award DBI#154689, NSF/EPSCoR Cooperative Agreement #IIA-1355423 and BioSNTR which is funded in part by the South Dakota Research and Innovation Center that supported this research.

## Supplementary information

**Figure S1.** Schematic representation of ‘Chambourcin’ experimental vineyard located at The University of Missouri Southwest Center Agricultural Experiment Station in Mount Vernon, Missouri, USA.

**Figure S2.** Complete ionomic results for 2014 and 2016 divided based on (A) rootstock (B) leaf position (C) rootstock by irrigation.

**Table S1.** Results for all factors explaining a significant portion of the variance for simple leaf shape descriptors consisting of aspect ratio, circularity, roundness and solidity. For each descriptor, the percent variance explained by the factor and the p-value are reported.

**Table S2.** Results for all factors explaining a significant portion of the variance for morphometric PC1 to 20. For each significant factor for a PC, the p-value, percent variance explained by the factor, and percent variance captured by the PC are all reported.

**Table S3.** Results for all factors explaining a significant portion of the variance for each element. For each significant factor for an element, the p-value and percent variance explained by the factor are reported.

**Table S4.** VitisNet Pathways that were uniquely positively enriched in a rootstock, or positively enriched in common for all three rootstocks, relative to own-rooted vines. A false discovery rate of 0.25 and nominal p-value of 0.05 were used to identify positive enrichment in each rootstock treatment.

**Table S5.** All genes which were significantly expressed in own-rooted vines were compared to genes in vines grafted to each rootstock to determine which ones were significantly differentially expressed. The results of these comparisons are listed. Annotations are from the VCost.v3 (Canaguier et al. 2017) reference annotation.

**Table S6.** Genes found to be significantly differentially expressed in vines grafted to only one rootstock when compared to own-rooted vines, or across vines grafted to all rootstocks compared to own-rooted vines, or not differentially expressed across any rootstock treatment, where tested for pathway enrichment.

