## Supplementary figures and images for "Rootstock effects on scion phenotypes in a ‘Chambourcin’ experimental vineyard"

### Supplementary file 1

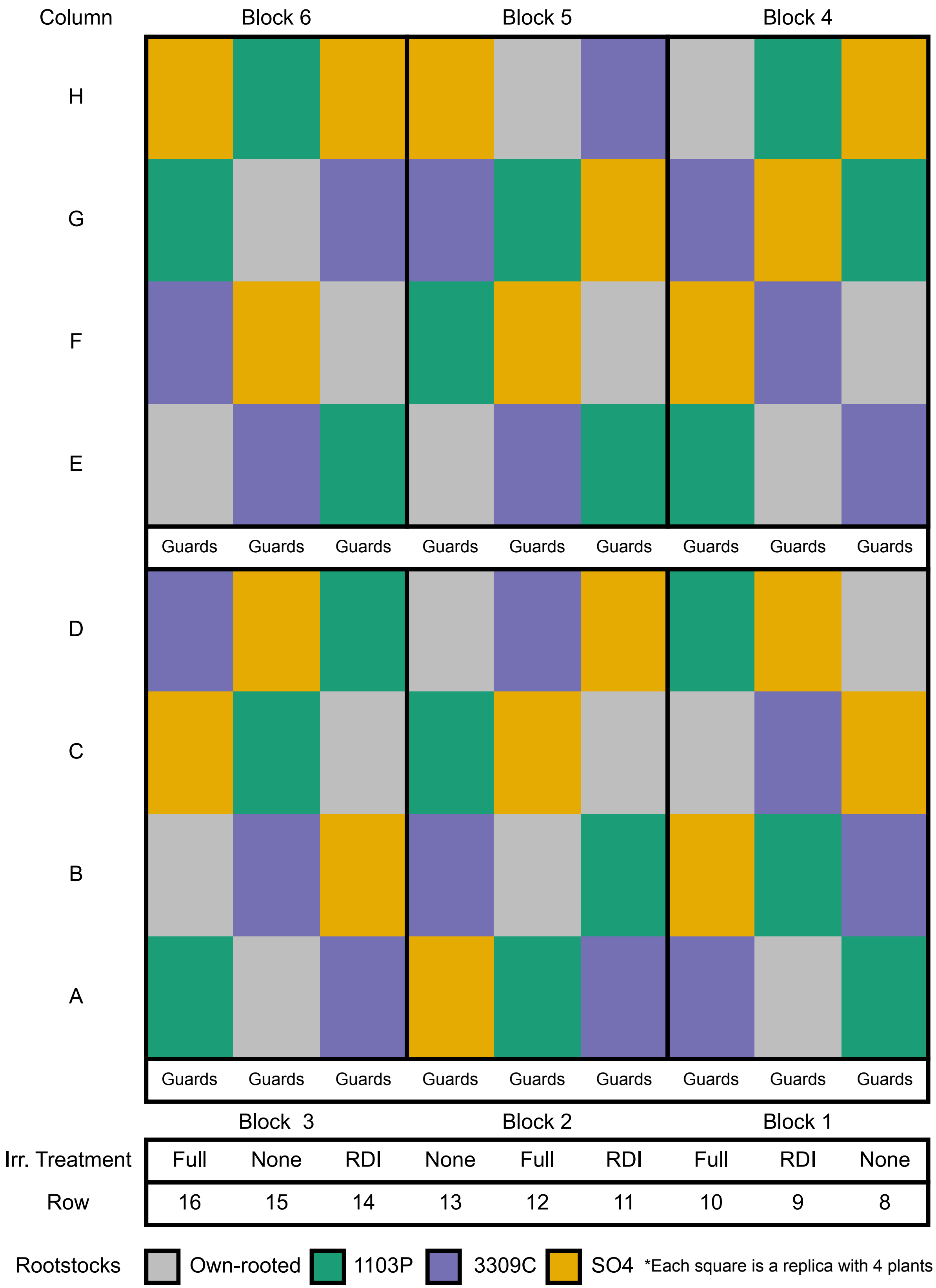

### Supplementary file 2

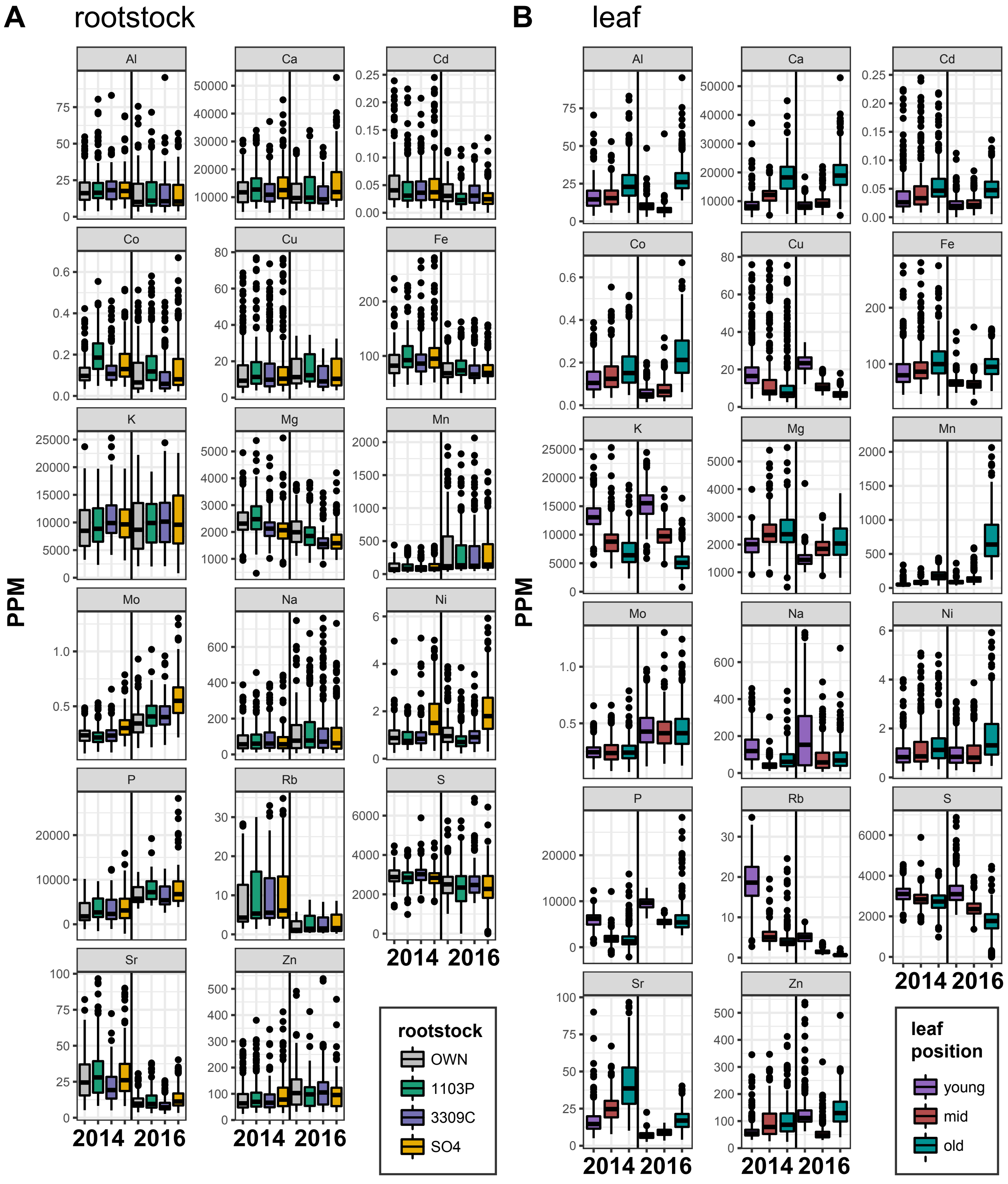

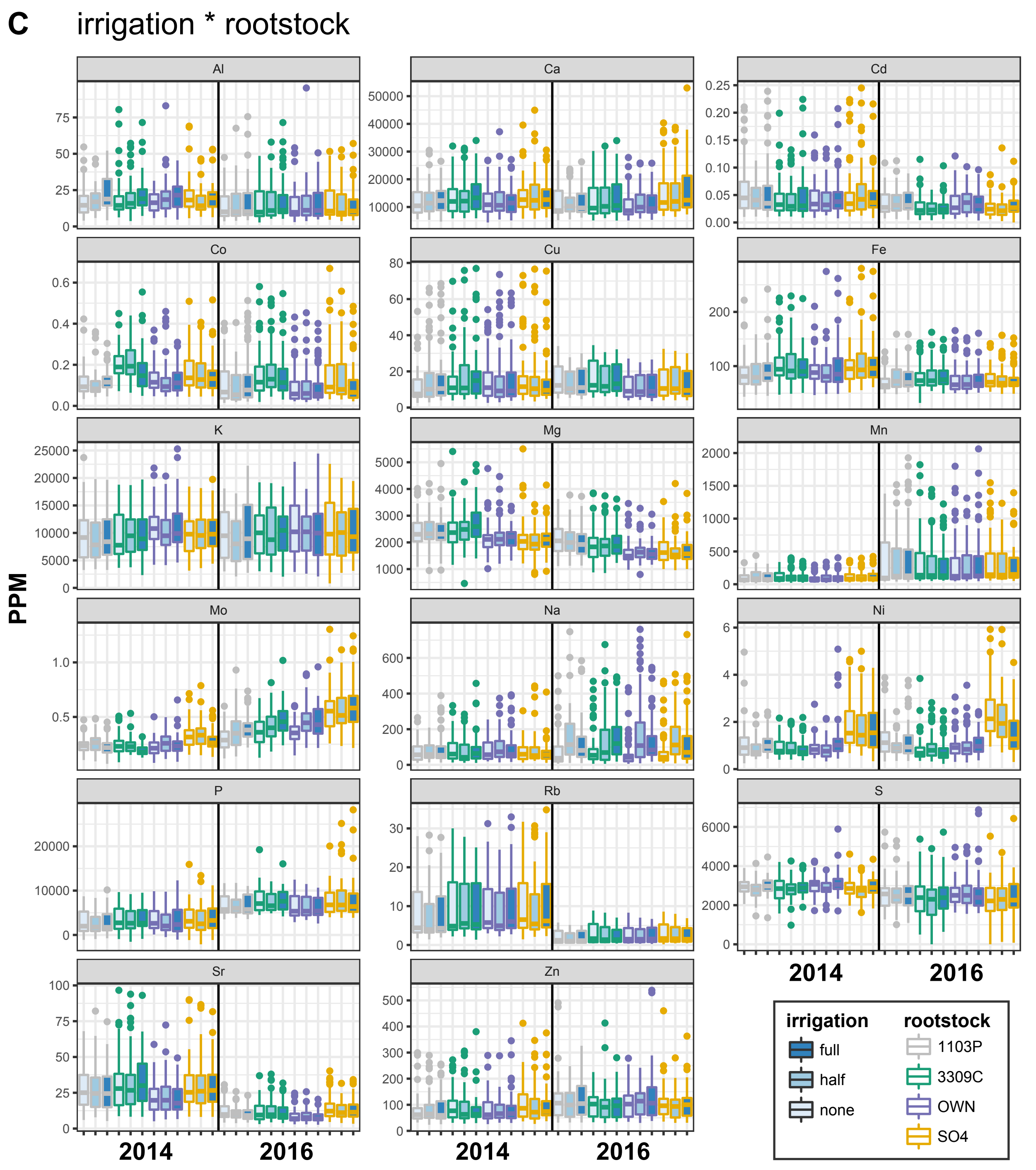
